# Genome-wide analyses of XRN1-sensitive targets in osteosarcoma cells identifies disease-relevant transcripts containing G-rich motifs

**DOI:** 10.1101/2020.08.11.246249

**Authors:** Amy L. Pashler, Benjamin P. Towler, Christopher I. Jones, Tom Burgess, Sarah F. Newbury

## Abstract

XRN1 is a highly conserved exoribonuclease which degrades uncapped RNAs in a 5’-3’ direction. Degradation of RNAs by XRN1 is important in many cellular and developmental processes and is relevant to human disease. Studies in *D. melanogaster* demonstrate that XRN1 can target specific RNAs, which have important consequences for developmental pathways. Osteosarcoma is a malignancy of the bone and accounts for 2% of all paediatric cancers worldwide. 5 year survival of patients has remained static since the 1970s and therefore furthering our molecular understanding of this disease is crucial. Previous work has shown a downregulation of XRN1 in osteosarcoma cells, however the transcripts regulated by XRN1 which might promote osteosarcoma remain elusive. Here, we confirm reduced levels of XRN1 in osteosarcoma cell lines and patient samples and identify XRN1-sensitive transcripts in human osteosarcoma cells. Using RNA-seq in XRN1-knockdown SAOS-2 cells, we show that 1178 genes are differentially regulated. Using a novel bioinformatic approach, we demonstrate that 134 transcripts show characteristics of direct post-transcriptional regulation by XRN1. Long non-coding RNAs (lncRNAs) are enriched in this group suggesting that XRN1 normally plays an important role in controlling lncRNA expression in these cells. Among potential lncRNAs targeted by XRN1 is *HOTAIR*, which is known to be upregulated in osteosarcoma and contribute to disease progression. We have also identified G-rich and GU motifs in post-transcriptionally regulated transcripts which appear to sensitise them to XRN1 degradation. Our results therefore provide significant insights into the specificity of XRN1 in human cells which is relevant to disease.

## INTRODUCTION

Spatial and temporal control of gene expression is critical to maintain cellular homeostasis. A crucial part of this regulatory network is the post-transcriptional control of RNA turnover in the cytoplasm. Deficiencies in RNA degradation can result in excesses of particular RNAs, which has implications for organism development, cell proliferation and a variety of human diseases including inflammation and viral infection (Astuti et al. 2012, Moon et al. 2015, Towler et al. 2015, Pashler et al. 2016, Towler et al. 2016, Towler and Newbury 2018, Towler et al. 2019). A major pathway operating within this network to provide post-transcriptional control of RNA expression is the 5’-3’ cytoplasmic RNA decay machinery. At the core of this pathway is the highly conserved 5’-3’ exoribonuclease XRN1. XRN1, the only cytoplasmic 5’-3’ exoribonuclease, functions as a complex with the decapping proteins DCP1/DCP2 (Braun et al. 2012) to remove the protective 5’ methylguanosine cap, resulting in an RNA with a 5’ phosphate which is susceptible to decay by XRN1.

Recent work suggests a model where XRN1-mediated decay is critical to maintain a complex regulatory feedback loop to control RNA Polymerase II (RNA pol II) activity (Abernathy et al. 2015, Gilbertson et al. 2018). Additional work has suggested that XRN1 itself is able to function as a transcriptional regulator in yeast cells (Blasco-Moreno et al. 2019). Modulation of XRN1 activity has been demonstrated to result in the cellular redistribution of a number of RNA binding proteins, which in turn affect RNA pol II activity (Gilbertson et al. 2018). XRN1 has also been demonstrated to be involved in co-translational decay (Tuck et al. 2020). Work in yeast has shown that XRN1 is able to directly interact with the ribosome, where the mRNA is directly channelled from the ribosomal decoding site into the active site of XRN1 (Tesina et al. 2019). Additionally, XRN1 has been shown to facilitate the clearance of transcripts on which the ribosome is stalled in mouse embryonic stem cells (Tuck et al. 2020). During nonsense mediated decay in mammalian cells, XRN1 rapidly removes the 3’ portion of the transcript after SMG-6-catalysed cleavage (Boehm et al. 2016). Therefore, XRN1 is plays a key role in many cellular pathways to regulate RNA levels.

Previous work in model organisms, such as *D. melanogaster, C. elegans* and *A. thaliana*, has shown null mutations/depletion of XRN1 results in specific developmental defects and/or lethality, strongly suggesting that XRN1 can target specific RNAs important in cellular or physiological processes. In *D. melanogaster*, null mutations result in defects during embryonic dorsal closure, small imaginal discs and lethality at the early pupal stage (Grima et al. 2008, Jones et al. 2012, Jones et al. 2013, Waldron et al. 2015, Jones et al. 2016). A key target in the larval stage is *dilp8*, encoding a secreted insulin-like peptide, which is known to co-ordinate developmental timing (Colombani et al. 2012, Jones et al. 2016). In *C. elegans*, knockdown of *xrn-1* results in defects in embryonic ventral enclosure and subsequent lethality, although the targets are unknown (Newbury and Woollard 2004). Whilst this work highlights the crucial developmental role of XRN1, the specific, physiologically relevant XRN1 targets in human cells remain elusive. The only well characterised role of XRN1 in human cells is during the host response to viral infection where its activity is inhibited, resulting in the stabilisation of short-lived RNAs such as FOS and TUT1 (Moon et al. 2012, Chapman et al. 2014, Moon et al. 2015).

Here we set out to identify and categorise XRN1-sensitive transcripts which are directly and indirectly sensitive to XRN1 activity in human cancer cells. Using modern techniques, we confirm previous findings by Zhang and colleagues (Zhang et al. 2002) to show that *XRN1* transcripts are reduced in levels in both osteosarcoma cell lines and patient samples, and extend these findings to two Ewing sarcoma cell lines. We identify a specific subset of transcripts that show sensitivity to XRN1 expression in osteosarcoma SAOS-2 cells and develop a method to assess the nature of their sensitivity. Using this method we demonstrate that transcripts that are directly and indirectly regulated by XRN1 are involved in specific cellular processes and display features which may confer their XRN1 sensitivity.

## RESULTS

### XRN1 is misexpressed in a subset of cancers of the mesenchymal lineage

XRN1 is an enzyme expressed ubiquitously with a critical role in regulating cytoplasmic RNA degradation. Semi-quantitative RT-PCR has been used previously to show that *XRN1* has reduced expression in human osteosarcoma cell lines and patient samples compared to foetal osteoblast (HOb) cells (Zhang et al. 2002). We confirmed these findings using modern quantitative PCR (qRT-PCR) on a range of human osteosarcoma cells lines and observed reductions in *XRN1* transcript levels in HOS and U-2 OS cells compared to HOb control cells. HOb cells were used as controls because they are primary foetal osteoblast cells and are not cancerous. No difference was observed in the SAOS-2 cell line, showing *XRN1* downregulation was not ubiquitous across osteosarcoma cell lines (Fig 1A). Interestingly, the HOS cell line, which expresses the lowest levels of *XRN1*, is also the most proliferative, whilst SAOS-2 cells, which do not show reduced *XRN1* expression proliferate more slowly (Sup Fig 1A). In contrast to *XRN1*, levels of other ribonucleases, *XRN2, DIS3, DIS3L1* and *DIS3L2* were not reduced, demonstrating that downregulation is specific to *XRN1* and not a general reduction in RNA stability mediators (Sup Fig 1B). Indeed, our results show an increase in the levels of all these other ribonucleases in HOS cells, suggesting a compensatory mechanism to maintain normal RNA levels. We then assessed the levels of *XRN1* pre-mRNA to test if transcription of *XRN1* was inhibited in these cells. Interestingly, we did not observe *pre-XRN1* downregulation in HOS or U-2 OS cells, suggesting the observed effects are a result of differential regulation at the post-transcriptional level (Fig 1B).

**Figure 1:**
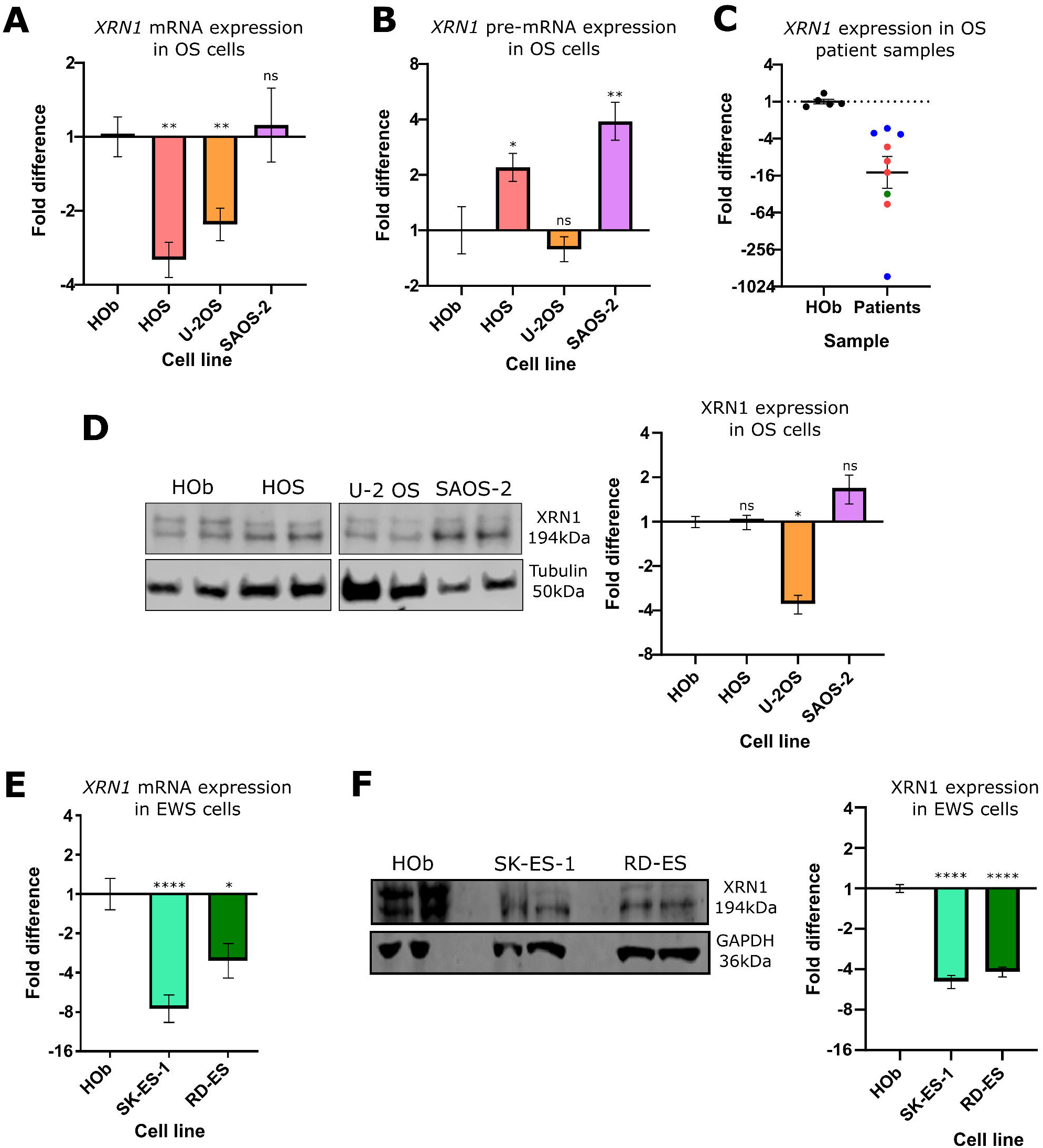
XRN1 is downregulated in osteo- and Ewing sarcoma. **A)** qRT-PCR quantification of *XRN1* mRNA expression across osteosarcoma (OS) cell lines in comparison to the HOb control cell line, normalised to *HPRT1*. Error bars represent SEM, n≥5. **B)** qRT-PCR quantification of *pre-XRN1* across osteosarcoma cell lines in comparison to the HOb control cell line, normalised to *HPRT1*. Error bars represent SEM, n≥6, **C)** qRT-PCR quantification of XRN1 across osteosarcoma patient samples in comparison to the HOb control cell line, normalised to *PES1*. Error bars represent SEM, n≥5, p=0.0296. Red = samples from hip and femur, blue=samples from scapula or humerus and green=unknown origin. **D)** Representative Western blot and graphical analysis showing expression of XRN1 protein in osteosarcoma cells as a proportion of that expressed in HOb control cells. Error bars represent SEM, n≥4. **E)** qRT-PCR quantification of *XRN1* mRNA expression in Ewing sarcoma (EWS) cell lines in comparison to the HOb control cell line, normalised to *GAPDH*. Error bars represent SEM, n≥6. **F)** Representative Western blot and graphical analysis showing expression of XRN1 protein in Ewing sarcoma cells as a proportion of that expressed in Hob control cells Error bars represent SEM, n≥4. For all figures ****=p<0.0001, **=p<0.01*=p<0.05 and ns=p>0.05.

To determine whether reduced levels of *XRN1* might have clinical importance in osteosarcoma we measured *XRN1* mRNA expression in 9 patient samples. Strikingly, all 9 samples showed reduced *XRN1* mRNA expression compared to HOb cells (Fig 1C). Western blotting confirmed the reduction in *XRN1* protein expression in U-2 OS cells, although a reduction in protein was not observed in HOS cells. Consistent with our qRT-PCR data, XRN1 protein expression was unaffected in SAOS-2 cells (Fig 1D). To test if our observations were specific to osteosarcoma progression, we next assessed XRN1 expression in the pathologically related bone sarcoma, Ewing Sarcoma. A decrease in both *XRN1* mRNA and protein was observed in two Ewing sarcoma cell lines, RD-ES and SK-ES-1, showing that our previous observations are not specific to osteosarcoma and suggesting XRN1 may have broader clinical importance (Fig 1E/F). Taken together these data demonstrate a need for further mechanistic understanding of the specific role played by XRN1 in these cells which could have clinical relevance.

### Phenotypic behaviour of SAOS-2 cells is not affected by XRN1 knockdown

Given the clear reduction of *XRN1* expression in the majority of osteosarcoma and Ewing sarcoma cells we set out to identify cellular processes specifically regulated by XRN1 within these cells. To achieve this, we performed a variety of phenotypic assays to determine the effect of XRN1 down regulation on cancer cell behaviour. For these experiments we used SAOS-2 cells as they showed wild-type levels of XRN1 expression compared to the HOb control. We hypothesised that depletion of XRN1 in SAOS-2 cells may induce a phenocopy of the HOS or U-2 OS cell lines which show an increased growth rate (Sup Fig 1A). Using siRNA we successfully reduced XRN1 expression to 20% of the levels observed in the scrambled siRNA controls within 24 hours. XRN1 protein levels remained depleted until at least 144hrs post transfection (Fig 2A and Sup Fig 2).

**Figure 2:**
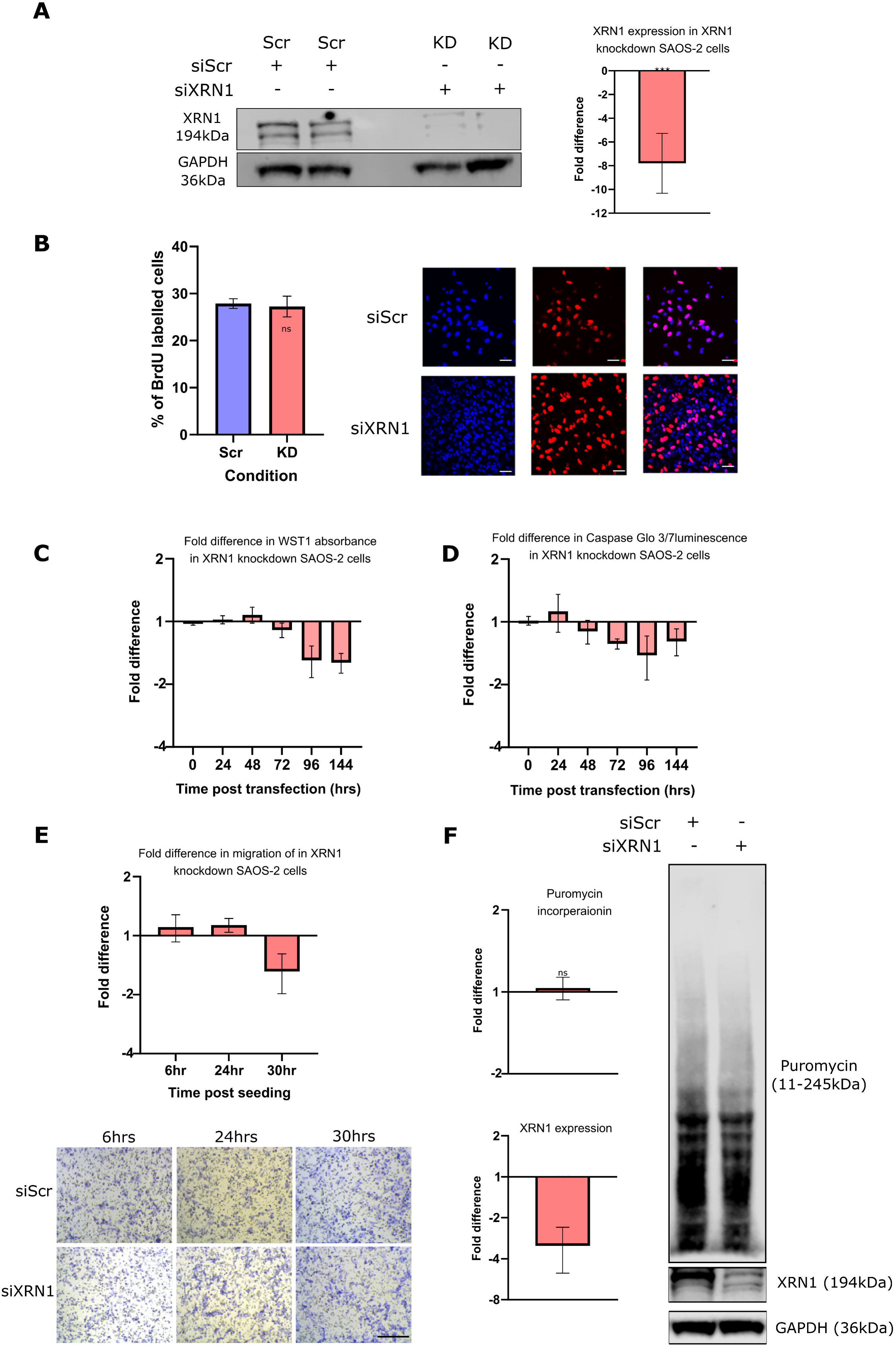
XRN1 knockdown in SAOS-2 cells does not result in observable phenotypes. **A)** Successful knockdown of XRN1 in SAOS-2 cells using RNAi 24 hours post transfection. Scr samples treated with 20pmol scrambled siRNA and KD cells treated with 20pmol *XRN1* siRNA. Error bars represent SEM, ***=p=0.0008. **B)** Quantification and representative images (40x objective) of the BrdU proliferation assay. Error bars represent SEM, n≥25, p=0.7938, scale bar=50µM. **C)** WST-1 assay at 24hr time intervals following transfection with either Scrambled (Scr) or *XRN1* (KD) siRNA. Error bar represent SEM, n=3. **D)** Caspase Glo 3/4 assay at 24hr time intervals following transfection with either Scrambled (Scr) or *XRN1* (KD) siRNA. Error bar represent SEM, n=3. **E)** Quantification and representative images (20x objective) of transwell migration assay 6hrs, 24hrs or 30hrs post seeding. Seeding was performed 24hrs post transfection with either Scrambled (Scr) or *XRN1* (KD) siRNA. Error bars represent SEM, n=4, p>0.05, scale bar=100µM. **F)** Knockdown of XRN1 does not affect nascent translation rates. Quantification of Puromycin incorporation or XRN1 expression (normalised to GAPDH relative to its own scrambled partner) 24hrs post transfection in cells treated with either Scrambled (Scr) or *XRN1* (KD) siRNA. Error bars represent SEM, ***=p=0.0003, ns=p=0.7432, n=5.

Using this model, we assessed proliferation and cell viability using BrdU staining and WST-1 assays, respectively. Although XRN1 expression was reduced by 81.8% we did not observe phenotypic changes when compared to the scrambled siRNA control (Fig 2B/C). Similarly, a Caspase-Glo 3/7 assay showed no strong change in the levels of apoptosis following XRN1 depletion (Fig 2D). In addition to viability and proliferation, cell migration is another crucial hallmark of cancer progression (Hanahan and Weinberg 2011). To assess if XRN1 depletion affects the rate of cell migration we used a transwell assay. However, we observed no changes in cell migration between XRN1-depleted and scrambled siRNA treated control cells over a 30-hour period (Fig 2E).

Finally, given that XRN1 has recently been shown to have strong roles in co-translational regulation in human and yeast cells (Tesina et al. 2019, Tuck et al. 2020) and translation factors are XRN4 targets in plant cells (Nagarajan et al. 2019) we hypothesised that the loss of XRN1 may affect translation rates. To test this we used SuNSET labelling to assess the rates of translation in XRN1-deficient cells. SuNSET labelling involves incubating cells with the tRNA analogue puromycin and subsequent blotting with a monoclonal α-puromycin antibody to detect and measure nascent translation. As puromycin is known to inhibit translation, careful optimisation of the concentration and time of incubation for each specific cell line was essential. We used 2.5µg/ml for 60 mins in SAOS-2 cells as we observed sufficient labelling whilst minimising the chances of saturation, in contrast to 10µg/ml which demonstrated reduced labelling after 60-90 mins, suggesting an inhibitory role on translation (Sup Fig 3). Although successful knockdown was achieved in each sample, we did not observe any difference in the rate of translation between XRN1 knockdown and scrambled siRNA control cells (Fig 2F). In summary, depletion of XRN1 in SAOS-2 cells does not appear to affect cell growth, viability, migration, or translation. It is possible, however, that XRN1 affects a phenotype we did not specifically test. Another possible reason is that immortalisation of SAOS-2 cells has been achieved through a mechanism not dependent upon XRN1, and that subsequent reduction in XRN1 level does not have an additive effect on this mechanism. Alternatively, there could be redundant or compensatory mechanisms within human cells following the loss of XRN1, although this seems unlikely based on observations in other organisms.

**Figure 3:**
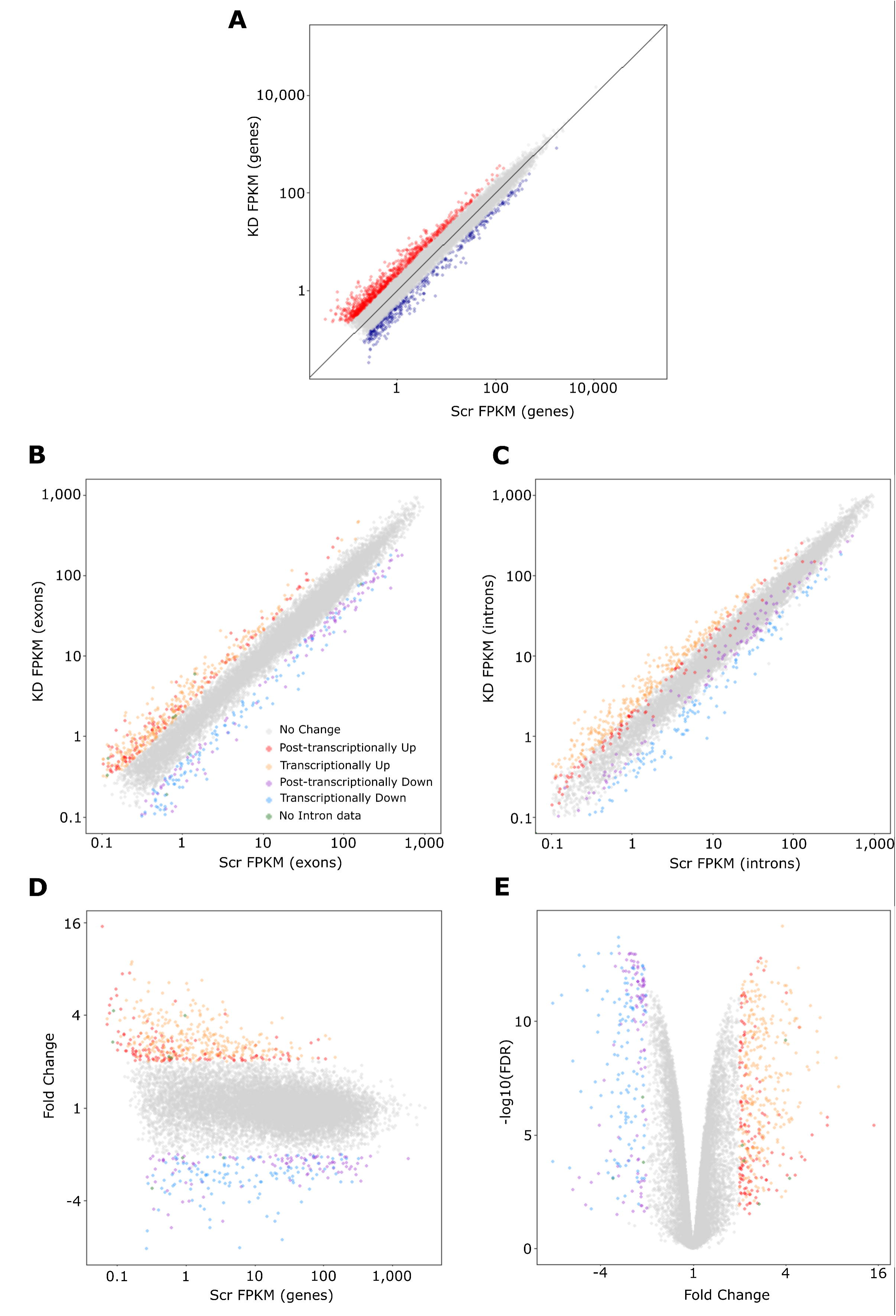
Overview of RNA-sequencing of XRN1-depleted SAOS-2 cells. **A)** Up- (red) and downregulated (blue) transcripts based on initial edgeR differential expression using genes as a counting method in featureCounts. **B)** Scrambled vs XRN1 knockdown FPKM based demonstrating differentially expressed transcripts using both gene and exon counting. Exon FPKM used for direct comparison with intron counting. Grey=no change, red=post-transcriptionally upregulated, orange=transcriptionally upregulated, purple=post-transcriptionally downregulated, blue=transcriptionally downregulated, green=no intron data. **C)** Differentially expressed transcripts when counting intron mapping reads allowing differentiation between transcriptional and post-transcriptional changes represented in B. Legend as in B. **D)** MA plot representing fold change in XRN1-depeleted SAOS-2 cells vs transcript expression in control cells coloured by nature of change. Legend as in B. **E)** Volcano plot demonstrating statistical information of all expressed transcripts. Legend as in B.

### RNA-sequencing reveals XRN1-sensitive transcripts in SAOS-2 cells

The results presented above show that although *XRN1* is post-transcriptionally depleted in human osteo- and Ewing-sarcoma cells and patient samples, its depletion appears to have no effect on the cell behaviours tested within SAOS-2 cells. We therefore adopted a molecular approach in order to identify transcripts that show specific sensitivity to XRN1 expression in SAOS-2 cells. By identifying these transcripts, we aimed to gain insights into the role of XRN1 in osteosarcoma cells.

We performed RNA-sequencing on SAOS-2 cells treated with either siRNAs to *XRN1* or a scrambled control, with 6 biological replicates for each condition; each XRN1 knockdown sample had a minimum *XRN1* depletion of 75% (Sup Fig 4A). For our initial analysis we removed adapters and quality trimmed raw RNA-sequencing files using Sickle and Scythe. Next, we used HiSat2 to map reads to the human genome (Ensembl release GRCh38.93). To account for potential expression changes due to conducting the knockdowns over consecutive weeks (1 Scrambled and 1 knockdown sample per week), we assessed gene expression using paired analysis. featureCounts was then used to count the number of reads mapping to each gene and paired differential expression analysis was subsequently performed using edgeR. Hierarchical clustering confirmed the paired nature of the samples, justifying our bioinformatic approach (Sup Fig 4B). Our analyses identified 1178 differentially expressed genes (defined as fold change >2 and FDR <0.05), of which 777 genes were upregulated and 401 genes were downregulated (Fig 3A). A greater number of upregulated transcripts is in line with the nature of XRN1 as an exoribonuclease with targets expected to increase in expression in the absence of XRN1.

**Figure 4:**
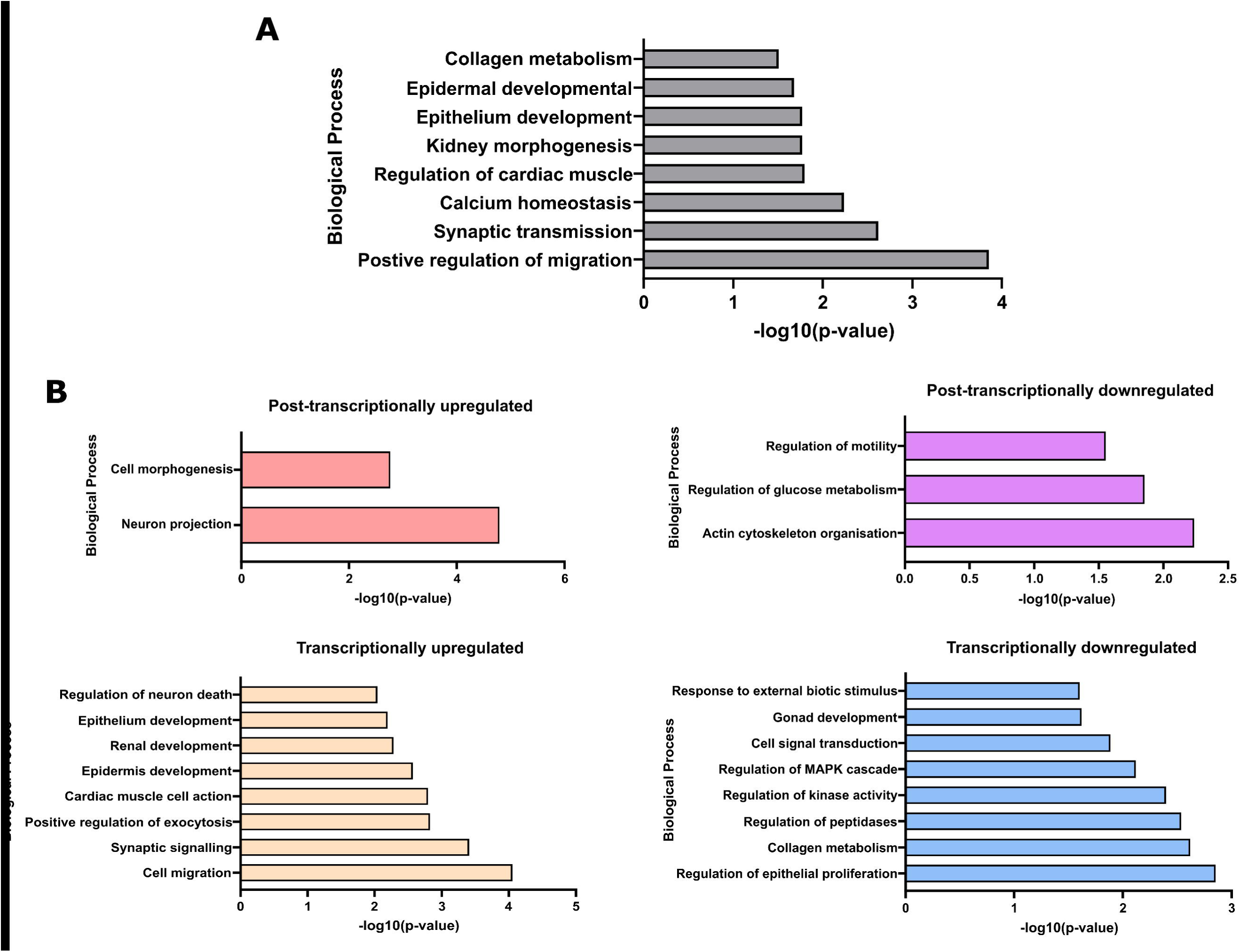
Gene Ontology analysis of differentially expressed transcripts. **A)** Gene ontology analysis using DAVID and Biological processes level “BPFAT” at highest stringency on all differentially expressed transcripts in XRN1-depleted SAOS-2 cells. **B)** As A, but enriched biological processes assessed in individual groups of misregulated transcripts.

While the initial analysis revealed a specific set of XRN1-sensitive transcripts, it did not explicitly identify those transcripts that are directly regulated by XRN1. For example, the 777 upregulated transcripts stabilised following XRN1 depletion, may represent direct effects (where transcripts are actively degraded by XRN1), or alternatively they could be transcriptionally upregulated as indirect consequences of loss of XRN1. We therefore re-purposed our analysis pipeline to allow genome wide assessment of transcriptional (indirect) and post-transcriptional (direct) effects of XRN1 depletion. To achieve this, we created a GTF annotation file containing the co-ordinates of every intron in the human genome. We then used featureCounts to count the number of exon (or intron) mapping reads in each XRN1 knockdown and control sample to find transcripts that increased post-transcriptionally. This was determined by identifying those with transcripts showing increases in exon-mapping reads but not in intron mapping-reads, indicating increased levels of mature mRNAs. Alternatively, those transcripts with increases in both exon and intron mapping reads would show increases in pre-mRNA, indicating increased transcription. The resulting count files were processed in a paired manner using edgeR and the same criteria were used to determine differential expression (fold change of >2 and an FDR of <0.05).

Using this approach, we saw high correlation between exon and gene related fold changes (Sup Fig 5, r^2^=0.91) with 722 transcripts passing the threshold in both samples (Fig 3B-E). When we included the intron level data, we observed a clear differentiation between post-transcriptional and transcriptional expression changes (Fig 3B/C). For example, transcriptionally upregulated transcripts (orange data points in Fig 3B/C) show increased expression at both the exon (Fig 3B) and intron (Fig 3C) levels. In contrast, 134 transcripts show the characteristics of post-transcriptional, direct regulation by XRN1 where increased expression is observed at the exon level but not the intron level (where the red data points in Fig 3C are within the grey, unchanged, region). We performed the same analyses on the downregulated transcripts and again observed examples of transcriptional (blue data points) and post-transcriptional (purple data points) changes in expression. We hypothesise that both transcriptional and post-transcriptional downregulation represent indirect effects due to XRN1 depletion. The transcripts that show post-transcriptional downregulation are likely to be themselves regulated by transcripts that are directly regulated by XRN1 such as miRNAs or those encoding RNA binding proteins. These analyses provide the first genome-wide differentiation between direct and indirect changes in gene expression following XRN1 depletion in human cells, summarised in Tables 1–4 and Supplemental File 1.

**Table 1:** Post-transcriptionally upregulated

| Ensembl GeneID | Gene name | Fold Change | FDR |
| --- | --- | --- | --- |
| ENSG00000237649 | KIFC1 | 15.02 | 3.96E-06 |
| ENSG00000108947 | EFNB3 | 7.51 | 3.85E-06 |
| ENSG00000183798 | EMILIN3 | 7.44 | 1.66E-06 |
| ENSG00000136274 | NACAD | 5.92 | 1.04E-05 |
| ENSG00000187867 | PALM3 | 5.34 | 0.000127488 |
| ENSG00000264569 | DCXR-DT | 5.07 | 9.66E-05 |
| ENSG00000186897 | C1QL4 | 4.94 | 1.79E-10 |
| ENSG00000197457 | STMN3 | 4.62 | 0.000610086 |
| ENSG00000246777 | AC044802.1 | 4.34 | 0.000109752 |

**Table 2:** Transcriptionally upregulated

| Ensembl GeneID | Gene name | Fold Change | FDR |
| --- | --- | --- | --- |
| ENSG00000164082 | GRM2 | 8.87 | 7.94E-08 |
| ENSG00000236609 | ZNF853 | 8.53 | 4.18E-09 |
| ENSG00000078900 | TP73 | 6.82 | 1.19E-06 |
| ENSG00000166341 | DCHS1 | 6.74 | 6.91E-11 |
| ENSG00000130592 | LSP1 | 6.50 | 2.33E-09 |
| ENSG00000172733 | PURG | 6.48 | 6.61E-09 |
| ENSG00000088881 | EBF4 | 6.38 | 2.70E-10 |
| ENSG00000141314 | RHBDL3 | 5.80 | 2.08E-11 |
| ENSG00000141750 | STAC2 | 5.69 | 1.42E-08 |

**Table 3:**
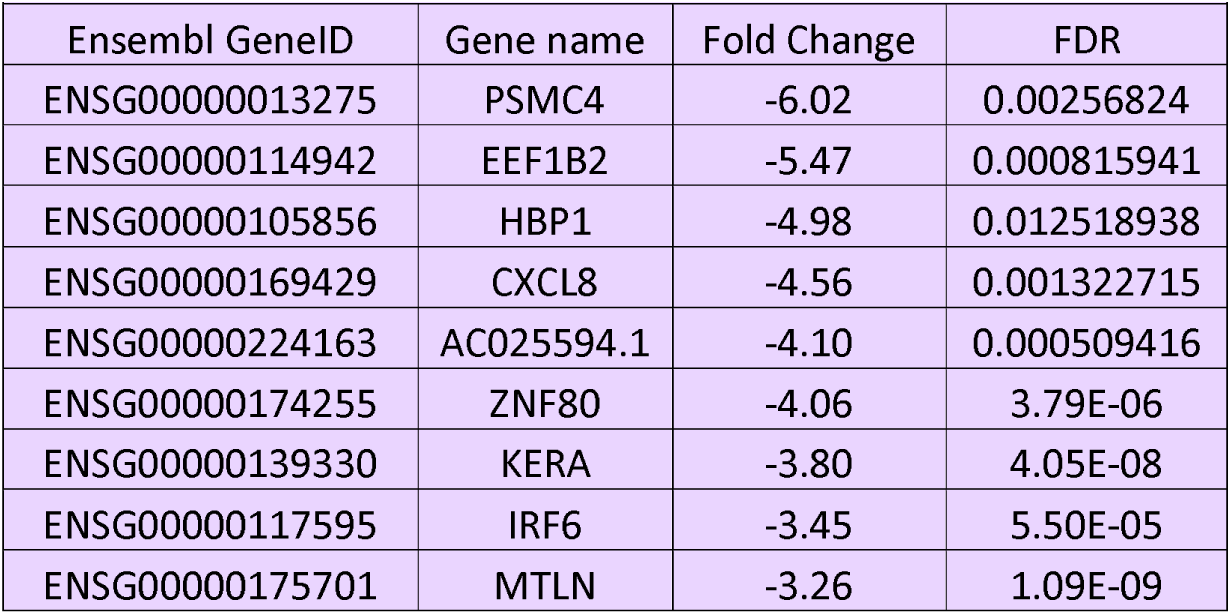
Post-transcriptionally downregulated

**Table 4:**
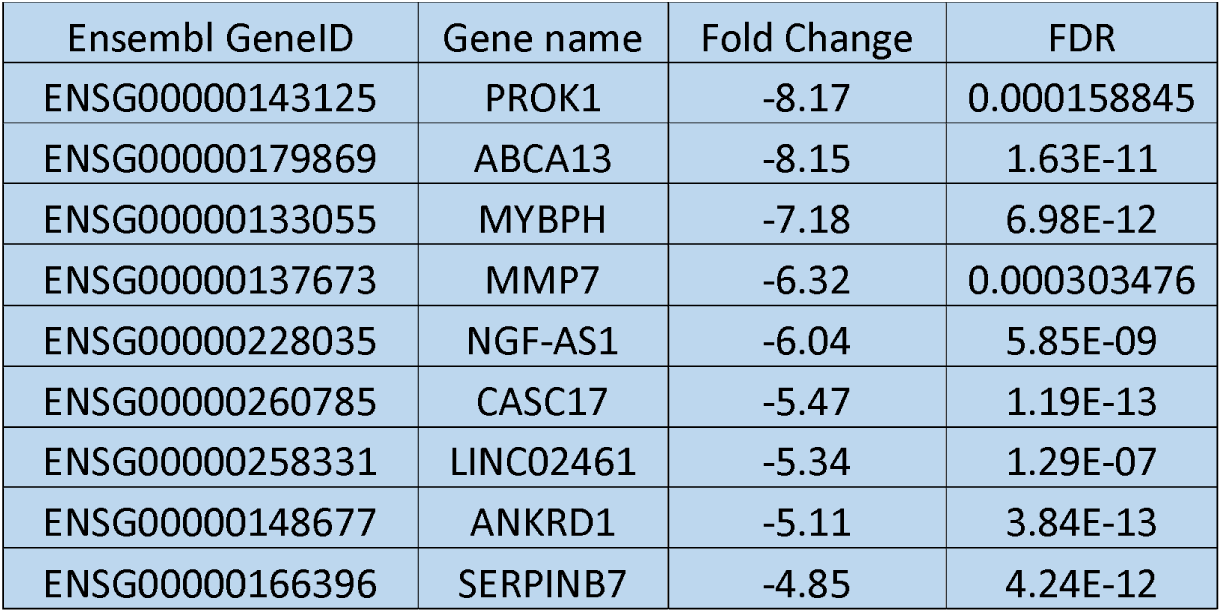
Transcriptionally downregulated

**Figure 5:**
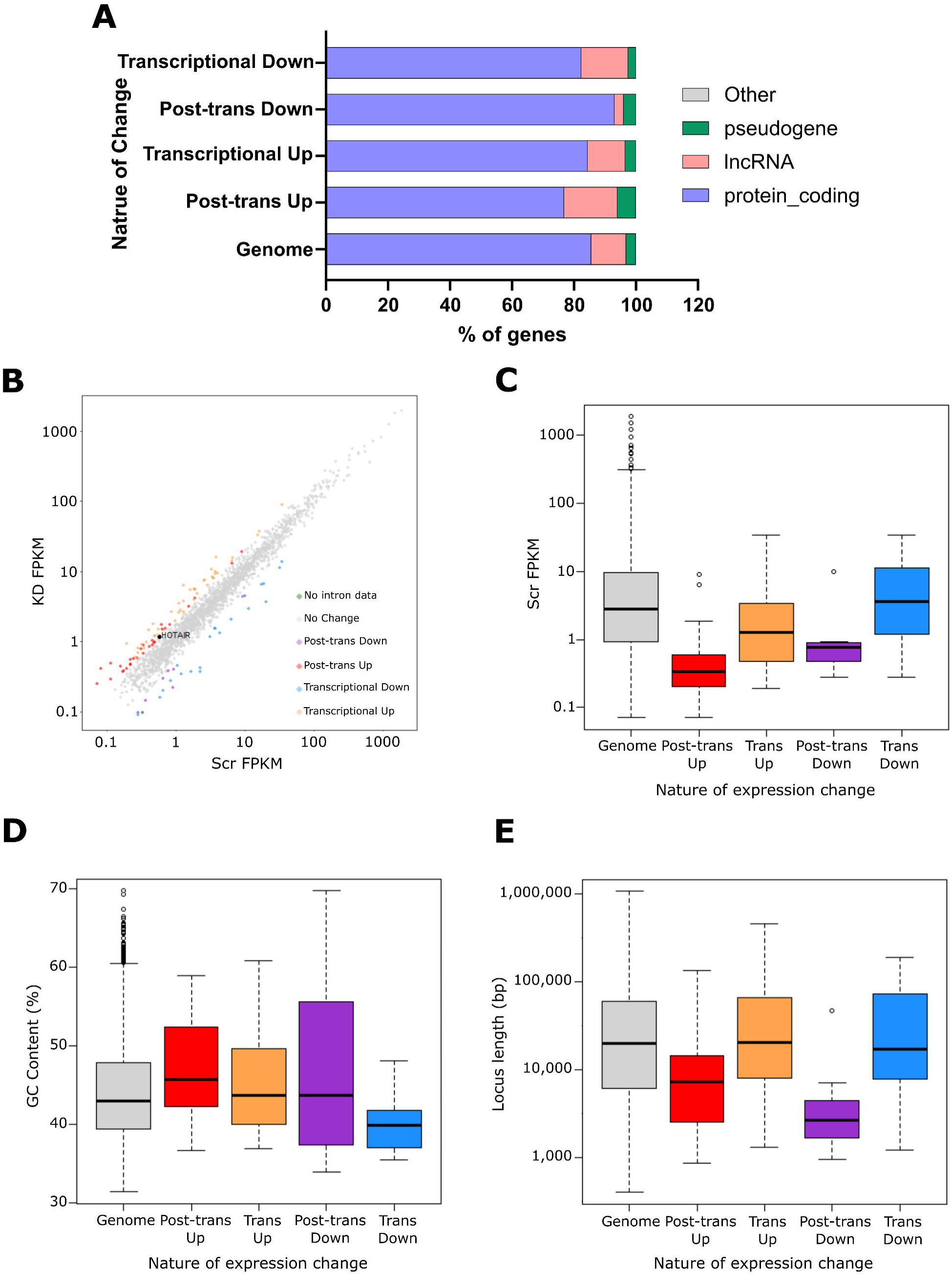
XRN1 also regulates ncRNAs in SAOS-2 cells. A) Assessment of transcript proportions affected by XRN1 depletion relative to the genome wide proportions detected in our sequencing data. *HOTAIR* highlighted in black, grey=no change, red=post-transcriptionally upregulated, orange=transcriptionally upregulated, purple=post-transcriptionally downregulated, blue=transcriptionally downregulated, green=no intron data. **B)** Scatter plot of changes in expression of all ncRNAs detected in our sequencing data. **C-E)** Boxplots of **C)** expression **D)** GC content and **E)** length (bp) of ncRNAs in our data. Grouped by their nature of change in expression and compared to the genome average as detected in our data set.

### XRN1-sensitive transcripts are involved in distinct biological processes

We used Gene Ontology (GO) analysis to identify specific biological processes affected following XRN1 depletion. Interestingly, GO analysis of all misregulated transcripts revealed an enrichment of genes involved in cell migration, a crucial hallmark of cancer progression (Fig 4A). However, we did not observe increased migration in our transwell assay, although this could be due to the nature of the knockdown experiments (discussed further below). Interestingly we also observe potential roles of XRN1 in epithelial and epidermal development. This is consistent with our previous work in *D. melanogaster* and *C. elegans* where we demonstrated that the XRN1 homologues are required for wound healing and epithelial sheet closure (Newbury and Woollard 2004, Grima et al. 2008). Movement of cell layers over other cells is also relevant for this solid cancer. We also observed a strong enrichment of transcripts involved in synaptic transmission suggesting a role for XRN1 in neuronal regulation. This is consistent with data showing that XRN1 forms discrete clusters associated with the post-synapse in hippocampal neurons and its knockdown impairs the translational repression triggered by NMDA (N-methyl-D-aspartate) (Luchelli et al. 2015).

To discriminate between the functional roles of transcripts directly and indirectly regulated by XRN1 we repeated our GO analysis with the specific sets of transcriptionally or post-transcriptionally up- or down-regulated transcripts (Fig 4B). This revealed that transcripts directly regulated by XRN1 have roles in cell morphogenesis and neurogenesis. Further, transcriptionally upregulated and post-transcriptionally down regulated genes are involved in a range of processes including epithelial development and cell migration. These analyses also demonstrate that transcriptionally downregulated genes are involved in cell signaling including the regulation of MAPK signaling.

### XRN1-sensitive transcripts demonstrate specific characteristics

Having identified transcriptional and post-transcriptional changes in gene expression following XRN1 knockdown in SAOS-2 cells, we wished to identify specific features or characteristics that may render the transcripts susceptible to XRN1-mediated decay. We first assessed the types of transcripts affected by loss of XRN1. A genome-wide assessment of transcript proportions detected in our samples revealed that 85.5% of detected RNAs were protein coding, 11.3% were lncRNAs, 3.1% were pseudogenes and the final <0.01% were classified as “other” transcripts (Fig 5A). Interestingly, whilst our transcriptionally up and down regulated groups mirrored the same proportions as the genome wide samples, ncRNAs appeared to be enriched amongst the post-transcriptionally upregulated genes with lncRNAs and pseudogenes representing 17.16% and 6% of the transcripts respectively (Fig 5A). Whilst the majority of misregulated transcripts were still protein coding (76.9%) this suggests that XRN1 directly regulates both mRNAs and ncRNAs in SAOS-2 cells. Of note is the post-transcriptional increase in expression of the lncRNA *HOTAIR* (2.11-fold, FDR<0.001) which is known to be upregulated in osteosarcoma cells and to contribute to disease progression (Wang et al. 2015, Li et al. 2017), suggesting a potential mechanistic link between XRN1-targets and osteosarcoma progression. Strikingly, lncRNAs were depleted from the post-transcriptionally downregulated transcripts (3% of the group). A possible explanation for this is that XRN1 normally targets miRNAs or transcripts encoding RNA binding proteins, which are then expressed at higher levels resulting in lower levels of their own target transcripts (Fig 5A).

Due to the enrichment of ncRNAs within the post-transcriptionally upregulated data set we next searched for features of these specific ncRNAs that may render them sensitive to XRN1-mediated decay. We first observed that the post-transcriptionally upregulated ncRNAs are usually expressed at low levels in control SAOS-2 cells (Fig 5B/C). We hypothesise that these ncRNAs are normally maintained at low levels of expression as a result of XRN1-mediated degradation. Interestingly, the post-transcriptionally regulated ncRNAs have a higher GC content than the genome average (Fig 5D, grey) or those that are transcriptionally regulated (Fig 5D orange/blue). We also observed a slight reduction in GC content in those transcripts that are transcriptionally downregulated (Fig 5D, blue). It is important to note that there are only 7 post-transcriptionally downregulated ncRNAs (Fig 5D, purple) and therefore this data must be interpreted with caution. Finally, ncRNAs that are post-transcriptionally regulated are much shorter than the genome average or those that are either up- or downregulated in a transcriptional manner (Fig 5E).

Next, we set out to assess if these transcript characteristics were specific to ncRNAs or if they were observed across all the transcripts post-transcriptionally regulated by XRN1. We observed the same pattern in expression levels and GC content that was previously observed specifically for the ncRNAs suggesting XRN1-sensitive transcripts are at low levels of expression in control cells and have a higher GC content than the genome average (discussed later) (Fig 6A/B). A collection of recent studies have shown that XRN1 is able to directly interact with the ribosome and that the level of translation can influence the stability of an mRNA transcript (Hanson et al. 2018, Tesina et al. 2019, Wu et al. 2019). To test if XRN1 targets have specific translational features we utilised published ribosome profiling data. As ribosome profiling data is not available for SAOS-2 cells, we used published data from an alternative osteosarcoma cell line, U-2 OS (Jang et al. 2015). This revealed that upregulated transcripts are usually translated in a less efficient manner than the genome average (Fig 6C).

**Figure 6:**
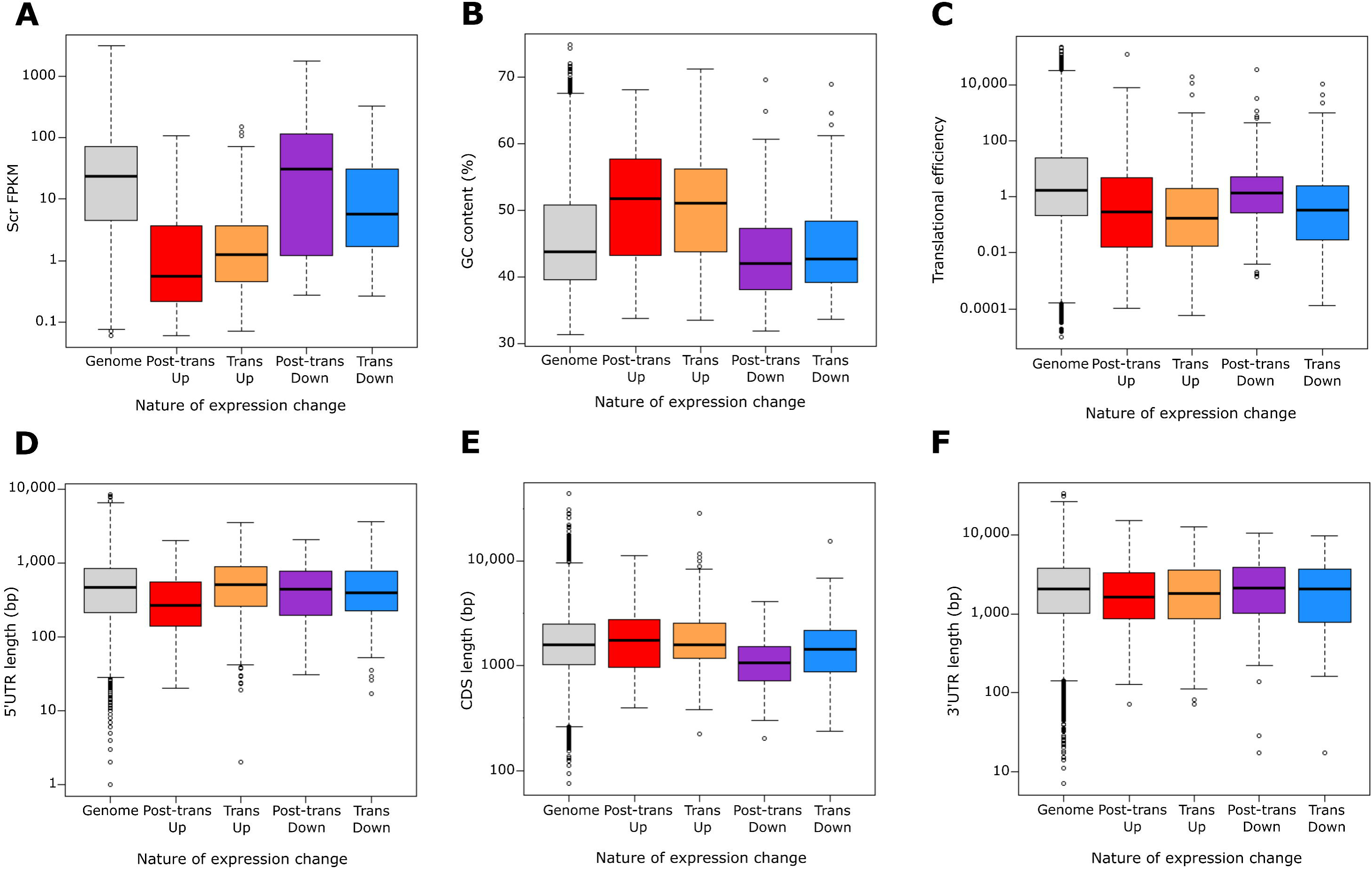
XRN1-sensitive transcripts show specific transcript characteristics. Boxplots of **A)** expression, **B)** GC content, **C)** translational efficiency, **D)** 5’UTR length, **E)** coding sequence (CDS) length and **F)** 3’UTR length of all transcripts within our data set. Grouped by their nature of change in expression and compared to the genome average as detected in our data set. Translational efficiency calculated as ribosome protected footprint FPKM/total RNA FPKM for each transcript.

**Figure 7:**
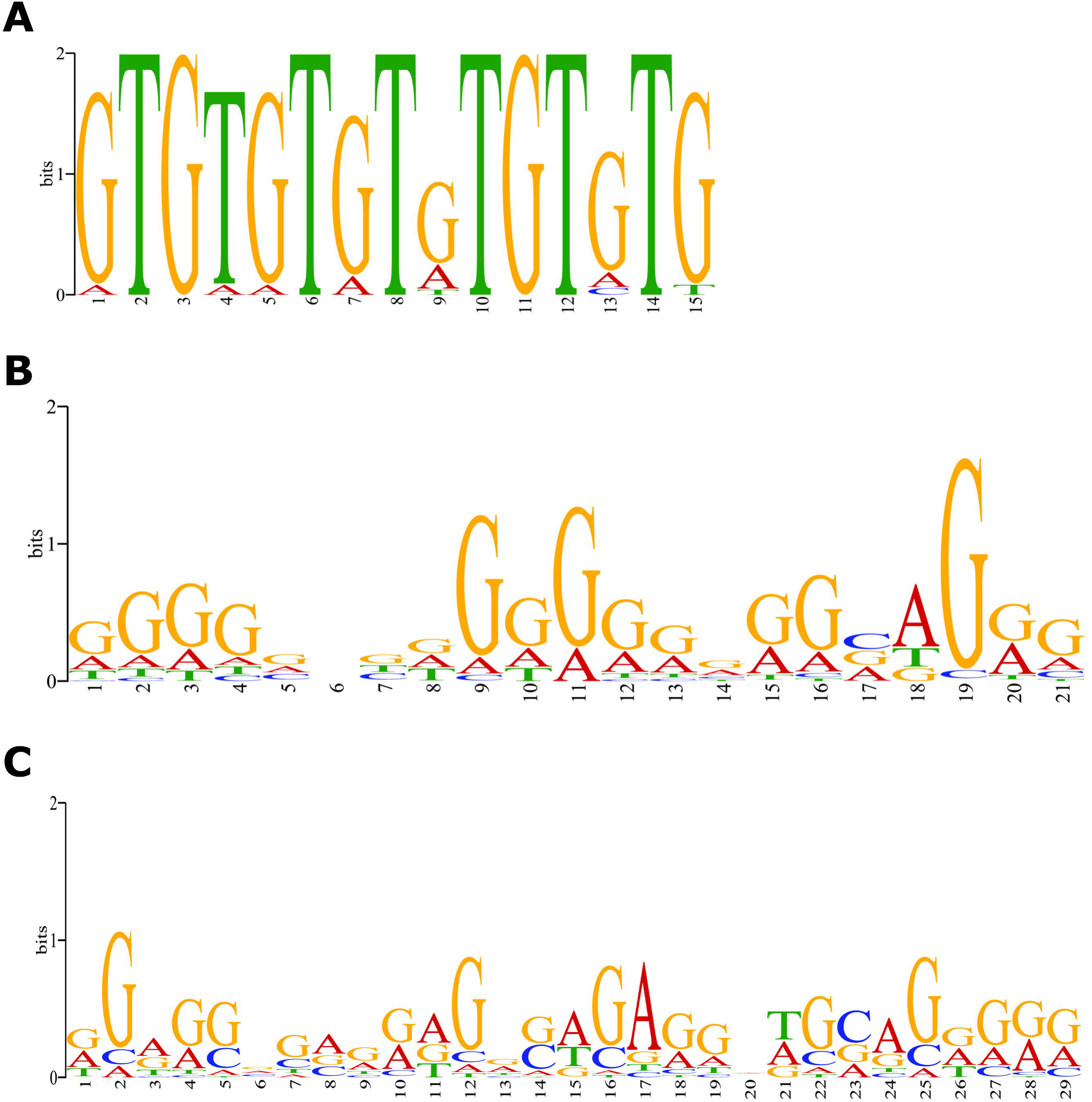
Direct XRN1 targets possess G-rich motifs. **A/B)** MEME analysis of 3’UTR of 103 mRNAs reveal **A)** GU rich 18 sites across 10 unique transcripts and **B)** G-rich 233 sites across 69 unique transcripts, motifs which may confer XRN1 sensitivity. **C)** Similar analysis of 30 ncRNAs post-transcriptionally upregulated in XRN1-depleted SAOS-2 cells reveals a similar G-rich motif to that observed in **B** (89 sites across 21 unique transcripts).

Finally, to specifically assess the features of mRNAs and compare with the previous ncRNA analyses we assessed the lengths of the major defined regions of an mRNA, the 5’ and 3’ Untranslated Regions (UTRs) and the coding sequence (CDS). This revealed that direct, post-transcriptional targets of XRN1 have shorter 5’UTRs than the genome average (272.5bp vs 401.1bp respectively, p<0.001) whilst the CDS was marginally longer and the 3’UTR was slightly shorter than the genome average (Fig 6D–F). Interestingly, the post-transcriptionally downregulated genes had a shorter CDS than the genome average, a phenomenon unique to this group of transcripts (1062.0bp vs 1600.9bp respectively, p<0.001) (Fig 6D–F). This suggests that these transcripts may have disproportionately long 3’ UTRs, which may render them susceptible to post-transcriptional regulators such as miRNAs and RNA binding proteins. Summary statistics for these analyses are shown in Tables 5/6.

**Table 5:** Average size of each mRNA region (bp)

| Table 5: Average size of each mRNA region (bp) |  |  |  |  |  |
| --- | --- | --- | --- | --- | --- |
| Region | Genome | Post-trans up | Trans up | Post-trans down | Trans-down |
| 5'UTR | 401.1 | 272.5 | 427.5 | 382.0 | 364.9 |
| CDS | 1600.9 | 1697 | 1732.3 | 1062.0 | 1414.6 |
| 3'UTR | 1819.3 | 1601.7 | 1713.5 | 1676.1 | 1648 |

**Table 6:**
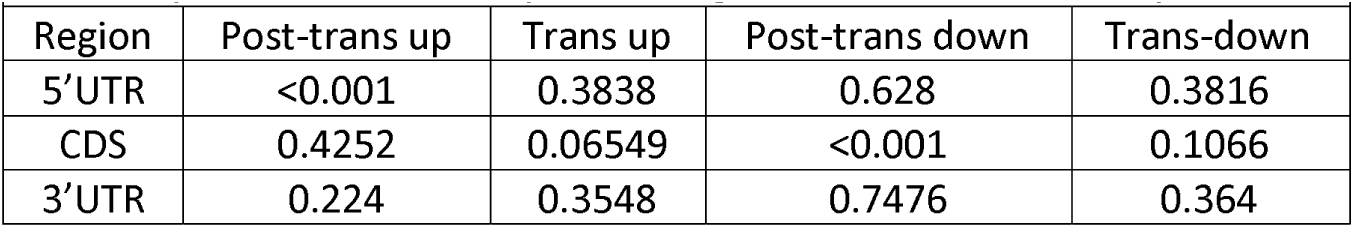
p-value of each comparison vs genome (Welch Two Sample t-test)

### Specific motifs may render transcripts susceptible to XRN1-mediated decay

mRNA 3’UTRs are known to control stability through *cis-acting* elements such as AU-rich elements (AREs). Therefore, we hypothesised that transcripts showing post-transcriptional upregulation (i.e. direct XRN1 sensitivity) may contain specific sequence motifs that allow for their targeting to XRN1 through interaction with other RNA binding proteins. To this end we used MEME (Bailey et al. 2009) to search the 3’UTR of 103 post-transcriptionally upregulated mRNAs for enriched motifs that may confer XRN1-sensitivity. This analysis revealed a section of significantly enriched motifs, of which 2 stood out; a G-rich motif (in 69/103 UTRs (67.0%)) and a second strong GU-rich motif in 10 (9.7%) of the 3’UTRs. Of the transcripts containing the GU-rich motif, all but one also include the G-rich motif (Supplemental File 2). Interestingly, GU-rich elements have been shown to function similar to AREs in promoting RNA decay so it is possible that GU-rich element binding proteins, such as the BRUNO/CELF family (Vlasova et al. 2008, Halees et al. 2011), may bind and promote 5’-3’ decay by XRN1. The most common motif shows a strong string of guanine residues which fulfill the criteria of forming G-quadruplexes. Recent work has shown that G-quadruplexes within 3’UTRs play important regulatory roles and consistent with the findings here, XRN1 has been shown to degrade transcripts containing G-rich regions more efficiently (Bashkirov et al. 1997).

Finally, as we also observed an enrichment of ncRNAs within the post-transcriptionally regulated transcripts, we performed a similar analysis using the whole ncRNA sequence to assess if similar motifs are identified. Analysis of the 30 post-transcriptionally regulated ncRNAs revealed a strikingly similar G-rich motif to that discussed above in 21 of the 30 submitted transcripts (70%). A total of 89 G-rich motifs were identified across these 21 transcripts with 6 sites within the ncRNA *HOTAIR*. These analyses suggest that this G-rich motif, which is likely to form G-quadruplex structures, is also able to sensitise specific transcripts to XRN1-medicated degradation in osteosarcoma cells. This novel finding suggests a new way that transcripts can be targeted for degradation by XRN1.

## DISCUSSION

Here we have expanded on previous findings using cell lines and patient samples to show that *XRN1* expression is reduced in osteosarcoma cells as well as in the cells of the related Ewing sarcoma. Using RNA-sequencing of XRN1 depleted SAOS-2 cells we performed a detailed genome-wide assessment of gene expression. We differentiated between transcriptional and post-transcriptional changes in expression and present a list of 134 transcripts that are likely to be direct targets of XRN1. Gene ontology analysis of differentially expressed transcripts revealed strong enrichment of transcripts associated with cell migration; a critical process required for cancer progression. This result is consistent with our previous findings in *D. melanogaster* and *C. elegans*, where depletion of Pacman or Xrn-1 result in defects in cell migration during embryonic dorsal closure and ventral enclosure respectively (Newbury and Woollard 2004, Grima et al. 2008). Transcripts directly regulated by XRN1 also appear to have roles in neurogenesis and neuron projection. Interestingly, proteins known to bind GU-rich regions, as identified in the MEME analysis have also been shown to be important regulators of neuronal gene regulation (Gallo and Spickett 2010, Dasgupta and Ladd 2012), and XRN1 activity may be important in the neurodegenerative disorder intranuclear inclusion body disease (Mori et al. 2018). XRN1 has also previously been shown to be localised in XRN1-positive bodies at the post-synapse in neurones where it contributes to local translational silencing elicited by NMDA (Luchelli et al. 2015).

Although RNA-sequencing revealed a number of transcripts that become misexpressed following loss of XRN1 in SAOS-2 cells, we observed no additional phenotypic defects within these cells. This is in contrast to XRN1 knockout HEK-293 cells which showed a 2-fold reduction in growth (Gilbertson et al. 2018). Although our RNA-sequencing experiments revealed differential expression of transcripts involved in regulating cell migration, migration rates over 30 hours were no different between XRN1-depleted and control cells. This could, however, be due to the use of RNA interference to deplete XRN1. Whilst we achieved a strong and consistent knockdown of ~80%, the 20% remaining may have sufficient residual activity to maintain cellular homeostasis. It is also possible that the changes in expression observed here were not sufficient in magnitude to elicit a phenotypic change. The lack of phenotype is intriguing given that deletion of the XRN1 homologue in *D. melanogaster*, Pacman, has severe phenotypic effects resulting in widespread apoptosis, reduction in tissue growth and male fertility, developmental delay and subsequent pupal lethality (Zabolotskaya et al. 2008, Jones et al. 2013, Waldron et al. 2015, Jones et al. 2016). The extensive conservation of XRN1 throughout eukaryotes suggests it has a critical function in maintaining homeostasis, however it is possible that in immortalised cell lines the role is less important. Another possibility is that SAOS-2 cells carry mutations that affect pathways redundant with XRN1 and therefore depletion of XRN1 may not present phenotypic effects. It is also conceivable that XRN1 in humans has a critical developmental role, as observed by the developmental phenotypes in *D. melanogaster* and *C. elegans* but these functions are specifically required in normal, multicellular tissues, rather than individual immortalised cells grown in culture.

We have identified specific sets of transcripts that are sensitive to XRN1 activity, including those directly regulated and those that are indirectly affected. We show that XRN1 is crucial for the direct regulation of both coding and noncoding RNAs, including the oncogenic lncRNA *HOTAIR*. Increased expression of *HOTAIR* has been shown to promote proliferation and metastasis of a variety of cancers (Özeş et al. 2016, Sharma Saha et al. 2016, Deng et al. 2017, Sun et al. 2017) and crucially has been frequently implicated in the progression of osteosarcoma (Wang et al. 2015, Li et al. 2017). Within these transcripts we identified specific motifs enriched in transcripts post-transcriptionally upregulated following XRN1 depletion, including a striking G-rich motif which is present in both mRNAs and ncRNAs directly regulated by XRN1. Recent work has shown that G-rich regions, that are capable of forming G-quadruplex structures, are crucial regulators of gene expression (Huppert et al. 2008). XRN1 shows increased efficiency of degrading transcripts containing G-rich regions (Bashkirov et al. 1997) and therefore it is possible that stretches of Guanine residues sensitise transcripts to XRN1-mediated decay, perhaps by the binding of particular RNA-binding proteins to GU rich regions, such as members of the CELF family, which in turn promote their decay via XRN1. Our results are also consistent with a previous study using HeLa and HCT116 cells, where transcripts with higher GC content are more sensitive to enzymes in the 5’-3’ degradation pathway such as DDX6 and XRN1 (Courel et al. 2019). The presence of this motif may also explain the increase in GC content in transcripts that show post-transcriptional upregulation. The ability of XRN1 to degrade G-rich RNAs is likely to be crucial as work on the cytoplasmic 3’-5’ ribonuclease Dis3L2 in *D. melanogaster* has revealed that Dis3L2 shows reduced efficiency for Guanine nucleotides and an absence of G-rich motifs within Dis3L2 targets (Reimão-Pinto et al. 2016, Towler et al. 2019). These transcripts may therefore normally depend on XRN1 for their degradation.

The data presented here also showed that XRN1 targets are normally maintained at low levels of expression and are likely to be rapidly turned over, similar to signatures of a number of oncogenes. Recent work in mouse embryonic stem cells has shown that XRN1 is directly recruited to the ribosome to remove transcripts that show reduced or stalled translation (Tuck et al. 2020). This is congruent with our findings that direct targets of XRN1 show reduced translational efficiency. It is possible that in the absence of XRN1, stalled/slowly translating ribosomes remain in contact with the RNA, increasing the chance of translational errors, which in turn could have detrimental effects upon the cell. Finally, XRN1-sensitive transcripts also tended to be shorter that the genome average, with shorter 5’UTRs. 5’UTRs are generally highly structured, therefore a shorter 5’UTR may result in a reduction of structure that would facilitate XRN1 activity. We also observe an array of indirect transcriptional changes in expression in XRN1-depleted SAOS-2 cells. This could be explained by recent work demonstrating that changes in expression or activity of XRN1 results in relocation of a number of RNA binding proteins. This includes other members of the decay machinery, which affect the mRNA-decay-RNA polymerase II transcriptional feedback loop (Abernathy et al. 2015, Gilbertson et al. 2018). Since an increase in XRN1 activity results in the relocation of a number of RNA binding proteins to the nucleus, it is possible that depletion of XRN1 causes these proteins to remain in the cytoplasm, contributing to the post-transcriptional downregulation of transcripts that we also observed here.

Taken together, the analyses presented in this study identify a number of features in coding and non-coding RNAs that may sensitise transcripts to XRN1-mediated decay. We present a group of high confidence direct targets of XRN1 in addition to a large group of transcripts that show indirect sensitivity to the ribonuclease. In the future, it would be of great interest to examine the identified motifs and features of these RNAs and begin to build a mechanism to explain the specificity of XRN1 targeting. This will shed light on the reasons for the selective downregulation of XRN1 in osteo- and Ewing-sarcoma cells.

## MATERIALS & METHODS

### Cell culture

Osteosarcoma cell lines, HOS, SAOS-2 and U-2 OS (ECACC), were cultured in DMEM-F12 (Gibco #21331-020) medium supplemented with 10% FBS (PAN-Biotech #P40-37100), 2mM L-Gln (Gibco #25030-024) and 100IU/mL penicillin, 100µg streptomycin (Gibco #15140-122). Cells were cultured at 37°C in a humidified incubator at 5% CO_2_. The foetal osteoblast cell line HOb (hFOB 1.19) (ECACC) was cultured in the same conditions. Ewing Sarcoma cell line, SK-ES-1 was cultured in McCoy’s 5A (Modified) medium (Gibco #26600-080) supplemented with 10% FBS, 2mM L-Gln and 100IU/mL penicillin, 100µg streptomycin. RD-ES was cultured in RPMI 1640 medium (Gibco #12633-020) supplemented in the same way. These Ewing sarcoma cell lines were provided by Prof. Sue Burchill, University of Leeds. Both were incubated at 37°C in a 5% CO_2_ humidified incubator.

### Patient samples

Samples were released by the Children’s Cancer and Leukaemia Group (CCLG) and sample details are outlined in Supplemental Table 1. Samples 11/650, 12/299 and 16/755 displayed large necrosis of the sample, and so were not included in analysis. Details of sample 16/591 were not disclosed.

### Western blotting

Western blots were performed on pellets of 1×10^6^ cells. Samples were run on 7% Tris-acetate Novex gels, apart from those used for SUnSET labelling, where samples were run on 4-12% Bis-Tris Novex gradient gels. GAPDH or Tubulin were used as loading controls. Blots were blocked in either 5% milk in 0.1% PBS-Tween or Odyssey Blocking Buffer (LI-COR #927-40000). Primary antibodies used were Mouse anti-GAPDH (1:10,000, Abcam #ab8245), Mouse anti-Tubulin (1:2000, Sigma #T9026) and Rabbit anti-XRN1 (1:2000, Bethyl Labs #A300-443A). Anti-mouse and anti-rabbit fluorescent antibodies were used at 1:20,000 (LICOR Donkey anti-mouse IR Dye 800CW and Goat anti-rabbit IRDye 680RD). Detection and quantification were performed using the LI-COR Odyssey Fc imager and Image Studio (version 5.2).

### qRT-PCR analysis

Total RNA was isolated from cell pellets and patient samples using a miRNeasy mini kit (Qiagen #217084) with on-column DNase digestion (Qiagen #79254). RNA concentrations were measured on a NanoDrop One spectrophotometer. Total RNA was converted to cDNA in duplicate using the High Capacity Reverse Transcription Kit (Applied Biosystems #4368814) and 500ng of RNA (according to manufacturer’s instructions) with random primers. A control ‘no RT’ reaction was performed in parallel to confirm that all genomic DNA had been degraded. qRT-PCRs were carried out on each cDNA replicate in duplicate (for a total of 4 technical replicates) using TaqMan Universal PCR Master Mix, No AmpErase UNG (Applied Biosystems #4324018) and TaqMan specific assays on a ViiA 7 or QuantStudio 7 machine. For the production of the custom *pre-XRN1* assay, the pre-mRNA sequence was submitted to Life Technologies’ web-based custom TaqMan Assay Design Tool as in (Jones et al. 2013). Standard TaqMan assays used in this study were to *XRN1* (ID:Hs00943063), *XRN2* (ID:Hs01082225), *DIS3* (ID:Hs0020014), *DIS3L1* (ID:Hs00370241) and *DIS3L2* (ID:Hs04966835). *GADPH* (ID:Hs02786624), *HPRT1* (ID:Hs02800695) or *PES1* (ID:Hs04963002) were used for normalisation.

### RNAi-mediated factor depletion

For siRNA transfections, 3×10^5^ SAOS-2 cells were seeded in a 6-well plate (34.8mm diameter). Transfections were carried out using Lipofectamine RNAiMAX reagent (Invitrogen #13778100) according to manufacturer’s instructions using Opti-MEM medium (Gibco #31985070) and DMEM-F12 medium without antibiotic. For each transfection, 20pmol of either siXRN1 (targeting exon 11, Invitrogen #125199) or siScrambled (Invitrogen #AM4611) were added for depletion of XRN1. For control cells 20pmol scrambled siRNA was added. siRNA was removed after 24 hours and replaced with fresh media.

### Phenotyping assays

Apoptosis assays were performed using Caspase-Glo 3/7 reagent according to manufacturer’s instructions (Promega #0000239042). Cell viability assays were performed using WST-1 cell viability reagent according to manufacturer’s instructions (Sigma #18993700). In both assays, 2×10^4^ cells were plated in black-walled 96-well plates overnight. XRN1 was knocked down using 5pmols siRNA in full medium and incubated for 24hrs. The reagent was then applied and luminescence (Caspase-Glow 3/7) or absorbance (WST-1) was measured on a plate reader. SUnSET labelling was performed using 2.5μg/mL puromycin (Merck #540411) incorporated into 4×10^5^ cells in 6-well plates where XRN1 had been knocked down for 24hrs. Puromycin was added for 1hr before cells were harvested and western blotting performed with GAPDH as a loading control, using an anti-puromycin antibody (Merck #MABE343). Puromycin incorporation was measured using Image Studio (version 5.2). Cell proliferation was determined by measuring Brd-U incorporation during DNA synthesis. Briefly, 10μM Brd-U (Sigma #B5002-100MG) was added to 5×10^4^ cells in a 24 well plate 24hrs post transfection with either siXRN1 or siScrambled (10pmols) for 6 hrs. Cells were subsequently fixed in 4% paraformaldehyde and permeabilised for 45 minutes in 0.3% Triton X-100 in PBS (PBTX). Following permeabilisation cells with incubated for 30 mins in 4M HCl follow by a 10-minute incubation in 0.1M sodium borate. Following washes in PBTX cells were incubated in αBrd-U diluted 1:20 in PBTX (Developmental Studies Hybridoma Bank G3G4). Cells were wased in PBTX before incubation in α-Mouse-Cy3 1:350 (Jackson ImmunoResearch #715-165-150). Cells were then washed in PBTX and mounted in Vectorshield containing DAPI (Vector Laboratories #H-1200). The ImageJ Dead_Easy Mitoglia Plug-In was used to measure the proportion of cells undergoing active DNA synthesis with the total number of cells counted using DAPI staining.

### RNA seq sample preparation and RNA library preparation

RNA was extracted from cell pellets, six replicates from siScrambled or siXRN1 treated cells were collected for sequencing over consecutive weeks. Total RNA was extracted using miRNEasy mini kit (Qiagen) with on-column DNase digestion (Qiagen). Total RNA concentration and quality were measured on a NanoDrop One, RNA integrity was assessed on an Agilent 2100 Bioanalyzer. RNA concentration was further assessed on a Qubit (Invitrogen). 500ng of total RNA was depleted for rRNA by Leeds Genomics using the Ribo-Zero kit. Library preparation was also performed by Leeds Genomics using the Illumina TruSeq standard protocol. Subsequent libraries were run in a 75bp single-end sequencing run on a Next Seq generating between 36 and 45 million reads per sample. Raw sequencing reads will be deposited in ArrayExpress following manuscript acceptance.

### Bioinformatic analysis of RNA-sequencing data

Sequence quality was assessed using FastQc c0.11.7 (http://www.bioinformatics.babraham.ac.uk/projects/fastqc/) and adapters were removed using Scythe v0.993b (https://github.com/vsbuffalo/scythe). Further quality control and read trimming was achieved using Sickle v1.29 (https://github.com/najoshi/sickle). The remaining high quality reads were mapped to the human genome GRCh38.93 from Ensembl using HiSat2 v2.01.0 (Kim et al. 2015) and SAM files were sorted and converted to BAM using SAMtools (Li et al. 2009). Paired analysis of control and knockdown cells was achieved using featureCounts (Liao et al. 2014) and edgeR (Robinson et al. 2010). Mapped reads were counted using featureCounts using GrCh38.93.gtf from Ensembl. Reads were counted at either the Gene, Exon or Intron level. For Intron data a novel.gtf file was computed from the exon boundaries within the original GRCh38.93.gtf. Only genes with a sum of 60 reads across the 12 biological replicates were retained for further analysis. Raw counts were used as an input for normalisation, quantification and differential expression analysis in edgeR. Transcripts were further filtered within edgeR and only those expressed in >10 samples were retained. Counts were then normalised and differential expression was assessed in a pairwise manner using the quasi likelihood F test where siXRN1 replicate 1 was compared to siScr replicate 1 and so on. Differentially expressed genes were initially determined as those showing a fold change of >2-fold and an FDR of <0.05. The same procedure was used for Exon and Intron level assessment, although intron reads were not filtered as this may have removed post-transcriptional changes. Post-transcriptional changes were determined as exon level changes of >2-fold and an FDR<0.05 and intron level changes of <2 fold or an FDR>0.05. The genes than showed changes at the exon and intron levels of >2 fold and an FDR<0.05 were classified as transcriptional changes.

### Data used that was not produced in this study

Translational efficiency data from U-2 OS cells was obtained from Jang et al 2015 where an average of all recorded time points was used. GC content and locus length were obtained from Ensembl using the BioMart tool.

### Gene Ontology and motif analysis

Functional annotation clustering of the differentially expressed genes was carried out using DAVID (Huang da et al. 2009, Huang da et al. 2009). Only the significantly enriched GO terms from the biological process category (BPFAT, using highest stringency and an enrichment score >1.3) were included in further analysis. 3’UTR and ncRNA motif analysis was conducted using the Meme Suite (Bailey et al. 2009).

### Statistical tests

All statistical analyses were performed in GraphPad Prism 8 or R (version 3.6.3). Unpaired Student t-tests were used to compare the means of single test groups to single control groups. Paired analysis with the quasi likelihood F test was used to determine differential gene expression using edgeR as outlined above. Welch’s two sample t-tests was used to determine significant changes in transcript features.

## ACKNOWLEDGEMENTS

The authors wish to thank Helen Stewart, Peter Bush, Sophie Robinson, Lisa Mullen and Stefano Caserta for helpful discussions. We would also like to thank Clare Rizzo-Singh for technical help and Sue Burchill (University of Leeds) for providing the Ewing sarcoma cell lines. Osteosarcoma samples were provided by the Children’s Cancer and Leukaemia Group (CCLG). This work was funded by a University of Brighton studentship [WC003-30] to A.L.P, a University of Brighton “Rising Stars Initiative” grant to C.I.J and S.F.N (WB002-34) and a Sussex Research Development grant to S.F.N and C.I.J (WC001-08). B.P.T was financed by a Biotechnology and Biological Sciences Research Council grant (BB/P021042/1) to S.F.N.

## AUTHOR CONTRIBUTIONS

A.L.P designed and performed most of the experiments and analysed some of the data. C.I.J. supervised and carried out the initial work, advised on the bioinformatics experiments and commented on the manuscript. T.B performed and analysed the experiments on Ewing sarcoma cells. B.P.T analysed and interpreted the RNA-seq data, prepared the Figures and wrote the majority of the manuscript. S.F.N. co-ordinated the study, contributed to the design and interpretation of the experiments and contributed to the writing of the manuscript.

---

**Supplemental Figure 1: Other ribonucleases are not downregulated in osteosarcoma cells. A)** Growth curves of HOS, U-2 OS and SAOS-2 cells. Error bars represent SEM, n=3. **B)** qRT-PCR assessment of *XRN2, DIS3, DIS3L1* and *DIS3L2* mRNA in osteosarcoma cell lines relative to the HOb control, normalised to *HPRT1*. Error bars represent SEM, n ≥3, ***=p<0.001, **p=<0.01, *=p<0.05, ns=p>0.05.

**Supplemental Figure 2: Time course of XRN1 knockdown in SAOS-2 cells.** Representative Western blot and quantification of all blots in cells treated with siScr (Scambled) or siXRN1 (KD) until 144 hours post transfection. Data presented relative to the paired scrambled control sample on each blot. Error bars represent SEM, n≥3, ***=p<0.001, **=p<0.01, *p<0.05.

**Supplemental Figure 3: Optimisation of SuNSET labelling experiments.** Error bars represent SEM (where n≥3). n≥2.

**Supplemental Figure 4: A)** qRT-PCR quantification of *XRN1* mRNA in each individual RNA-sequencing replicate. Each *XRN1* knockdown replicate shown in pink is compared to its paired scrambled control (Scr) replicate. Mean and SEM shown. **B)** Hierarchical clustering of RNA-sequencing samples following edgeR analysis.

**Supplemental Figure 5:** Correlation between transcript fold changes between edgeR analyses when either counting at the “gene” or “exon” level with featureCounts. r^2^=0.91.

